# Selective epithelial expression of *KRAS*^G12D^ mutation drives ductal pancreatic neoplasia in Oncopigs

**DOI:** 10.1101/2024.02.01.578513

**Authors:** Carlos P Jara, Al-Murtadha Al-Gahmi, Audrey Lazenby, Michael A. Hollingsworth, Mark A. Carlson

## Abstract

**Background:** Pancreatic ductal adenocarcinoma (PDAC) remains a formidable challenge in oncology, characterized by a high mortality rate, largely attributable to delayed diagnosis and the intricacies of its tumor microenvironment. Innovations in modeling pancreatic epithelial transformation provide valuable insights into the pathogenesis and potential therapeutic strategies for PDAC.

**Methods:** We employed a porcine (Oncopig) model, utilizing the Ad-K8-Cre adenoviral vector, to investigate the effects of variable doses (10^7^ to 10^10^ pfu) on pancreatic epithelial cells. This vector, the expression from which being driven by a Keratin-8 promoter, will deliver Cre-recombinase specifically to epithelial cells. Intraductal pancreatic injections in transgenic Oncopigs (*LSL*-*KRAS*^G12D^-*TP53*^R167H^) were performed with histologically based evaluation at 2 months post-injection.

**Results:** Specificity of the adenoviral vector was validated through Keratin-8 expression and Cre-recombinase activity. We confirmed that the Ad-K8-Cre adenoviral vector predominantly targets ductal epithelial cells lining both large and small pancreatic ducts, as evidenced by Keratin 8 and CAM5.2 staining. Higher doses resulted in significant tissue morphology changes, including atrophy, and enlarged lymph nodes. Microscopic examination revealed concentration-dependent proliferation of the ductal epithelium, cellular atypia, metaplasia, and stromal alterations. Transgene expression was confirmed with immunohistochemistry. Desmoplastic responses were evident through vimentin, α-SMA, and Masson’s trichrome staining, indicating progressive collagen deposition, particularly at the higher vector doses.

**Conclusion:** Our study suggests a distinct dose-response relationship of Ad-K8-Cre in inducing pancreatic epithelial proliferation and possible neoplasia in an Oncopig model. All doses of the vector induced epithelial proliferation; the higher doses also produced stromal alterations, metaplasia, and possible neoplastic transformation. These findings highlight the potential for site-specific activation of oncogenes in large animal models of epithelial tumors, with the ability to induce stromal alterations reminiscent of human PDAC.

## INTRODUCTION

Pancreatic ductal adenocarcinoma (PDAC) remains a challenge in public health, responsible for 7% of all cancer-related deaths. Five-year survival rate is approximately 9%, reflecting the disease’s aggressive nature and modestly effective treatment options (1-3). The use of large animal models, particularly porcine models, may provide a supplemental and unique approach to PDAC research, especially with respect to porcine physiological and anatomical similarities to humans (4). Porcine size enables the replication of human interventional procedures which are challenging to perform in rodent models. Furthermore, the porcine immune system, lifespan, and genetic sequence mirrors those of humans better than mice do (4, 5).

In a previous study (6), we employed the transgenic Oncopig model (somatic *LSL*-*KRAS*^G12D^-*TP53*^R167H^) (7) to describe and characterize a porcine model of pancreatic cancer (proof-of principle for such a model was published earlier) (8, 9). The KRAS^G12D^ mutation is a near-ubiquitous initiator and driver in PDAC (10-15). In the Oncopig we used pancreatic ductal injection of an adenovirus that expressed Cre-Recombinase under the control of CMV promoter, allowing high Cre expression in almost any cell type, resulting in expression of the two mutants (KRAS^G12D^ and p53^R167H^) in infected cells. The outcome was fulminant tumor growth which led to subject demise in 2-3 weeks; tumor pathology was mixed with an epithelial component (6).

In order to have a less fulminant, more tractable porcine pancreatic cancer model which also would be fully epithelial in origin, we utilized in the present study an adenovirus which would express Cre-Recombinase under control of the Keratin 8 promoter, a vector whose Cre expression would be restricted to epithelium (16). We believed that the degree of transformation observed in response to varied adenoviral vector doses might provide an improved porcine model of pancreatic cancer that could be used for device development, testing of antitumor drugs, or even study of early tumor pathogenesis and tissue response.

## RESULTS

### Keratin-8 And Cre-Recombinase Expression In Pancreatic Cells

First, we explored Keratin-8 expression in pancreatic epithelial cells by using single-cell RNA sequencing from whole human pancreas. We successfully identified higher levels of Keratin-8 RNA expression mainly in pancreatic ductal cells and lower in acinar cells, both in human healthy and PDAC samples (Fig. 1a). Next, we explored Keratin 8 protein expression in the porcine pancreas using immunohistochemistry (IHC). We identified epithelial staining (CAM5.2 staining and Keratin 8) in accessory and main ducts in both wild type and Oncopig pancreas (Fig. 1b). Despite human scRNA sequencing results indicating some K8 expression in acinar cells, our analysis in porcine pancreas showed Keratin-8 staining was restricted primarily to ductal structures, with no detectable expression in acinar cells. Finally, we performed an e*x vivo* assessment of the Ad-K8-Cre Adenovirus (a vector construct with Keratin 8 promoter driving the expression of the Cre-Recombinase) in wildtype porcine pancreas (Fig. 1c, Supplementary S1a). This *ex vivo* infection assay showed that the expression of Cre-recombinase under the control of the Keratin 8 promoter was limited to the epithelial cells, confirming the selective activity of the adenoviral construct in pancreatic epithelial cells (Fig. 1d).

**Figure 1:**
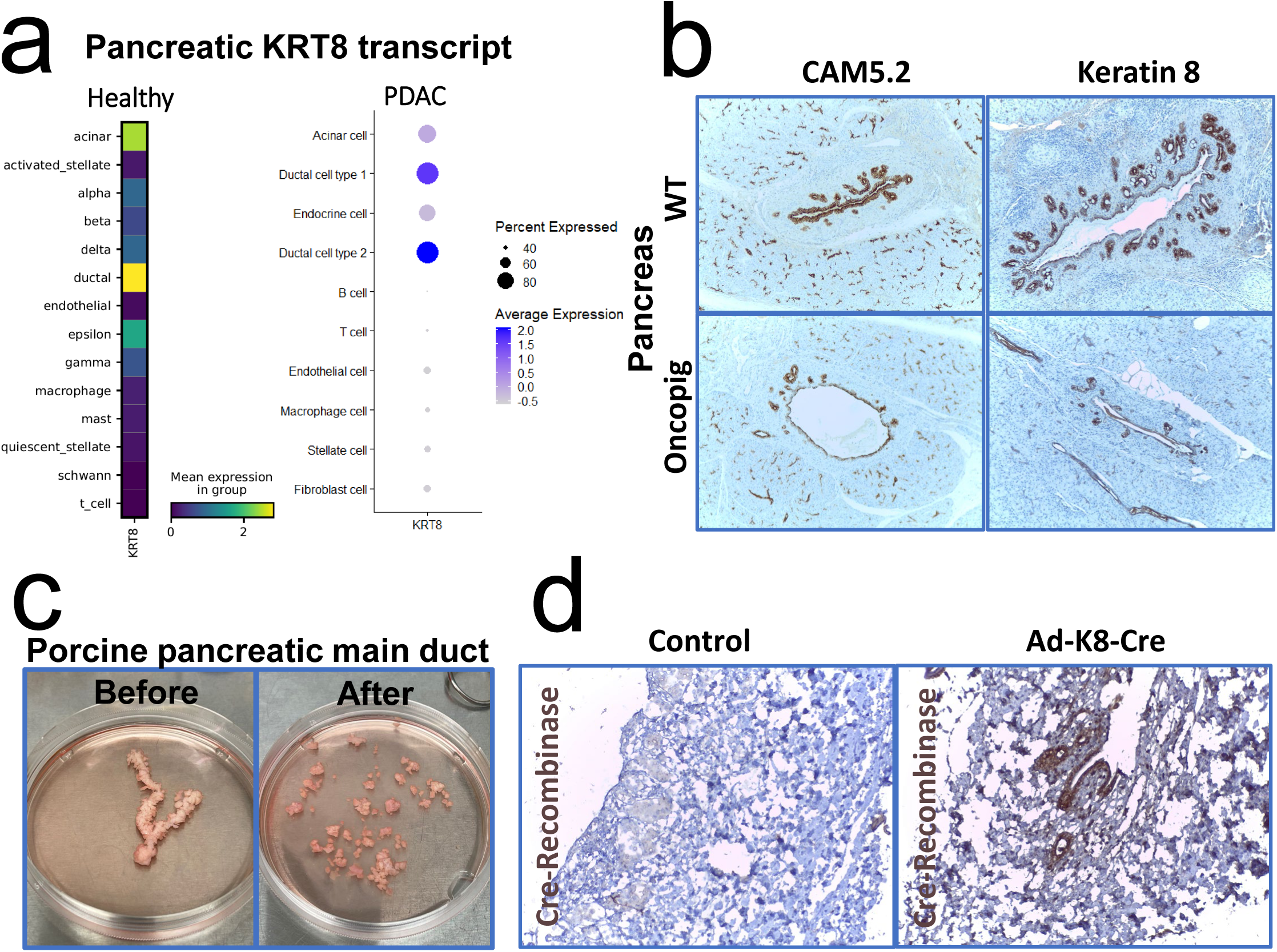
Analysis of Keratin 8 Expression and Activity in Pancreatic Cells. (a) Single-Cell RNA Sequencing Analysis of KRT8 in Pancreatic Cells: Heatmap (left) displays mean expression of KRT8 transcript across various cell types in healthy pancreas. Dot plot (right) highlights the percentage of KRT8-positive cells and average expression levels in cell types from pancreatic ductal adenocarcinoma (PDAC) samples. Data derived from 26 pancreatic cancer and 13 control pancreas samples. (b) Immunohistochemical Detection of Keratin 8 in Pancreatic Sections: Representative staining images of CAM5.2 and Keratin 8 in pancreatic tissue. Top row: Wild type (WT) pancreas; Bottom row: Oncopig pancreas. Staining emphasizes Keratin 8 presence in both primary and accessory pancreatic ducts. (c) Ex Vivo Analysis of Adenoviral Activity in Porcine Pancreatic Duct: Before (left) and after (right) images of porcine pancreatic main duct tissue preparation. This ex vivo assay assesses adenovirus-mediated Cre-recombinase activity driven by the Keratin 8 promoter, confirming selective targeting of pancreatic ductal cells. (d) Cre-Recombinase Expression Following Ad-K8-Cre Administration: Immunohistochemistry for Cre-recombinase in pancreatic sections. Control (left) and Ad-K8-Cre-treated (right) samples show targeted Cre expression in ductal epithelial cells, verifying the specificity of the Keratin 8 promoter in driving gene expression in pancreatic ducts.

### Dose-Dependent Macroscopic Alteration After Vector Administration

After confirming targeted expression of Cre-recombinase by Ad-K8-Cre in pancreatic epithelial cells, we evaluated for targeted epithelial activity of the Ad-K8-Cre adenovirus *in vivo* (Supplementary S1b). We injected Ad-K8-Cre adenovirus into the duct of the connecting lobe of the pancreas, as previously described (6), at different concentrations (ranging from 10^7^ to 10^10^ pfu) in eight transgenic Oncopigs (Supplementary S2a-c). For the purposes of this report, the injection region, or the portion of pancreas subjected to the vector treatment, is meant to be synonymous to with the region of the connecting lobe of the pancreas (dashed oval in Fig. 2a, b).

**Figure 2:**
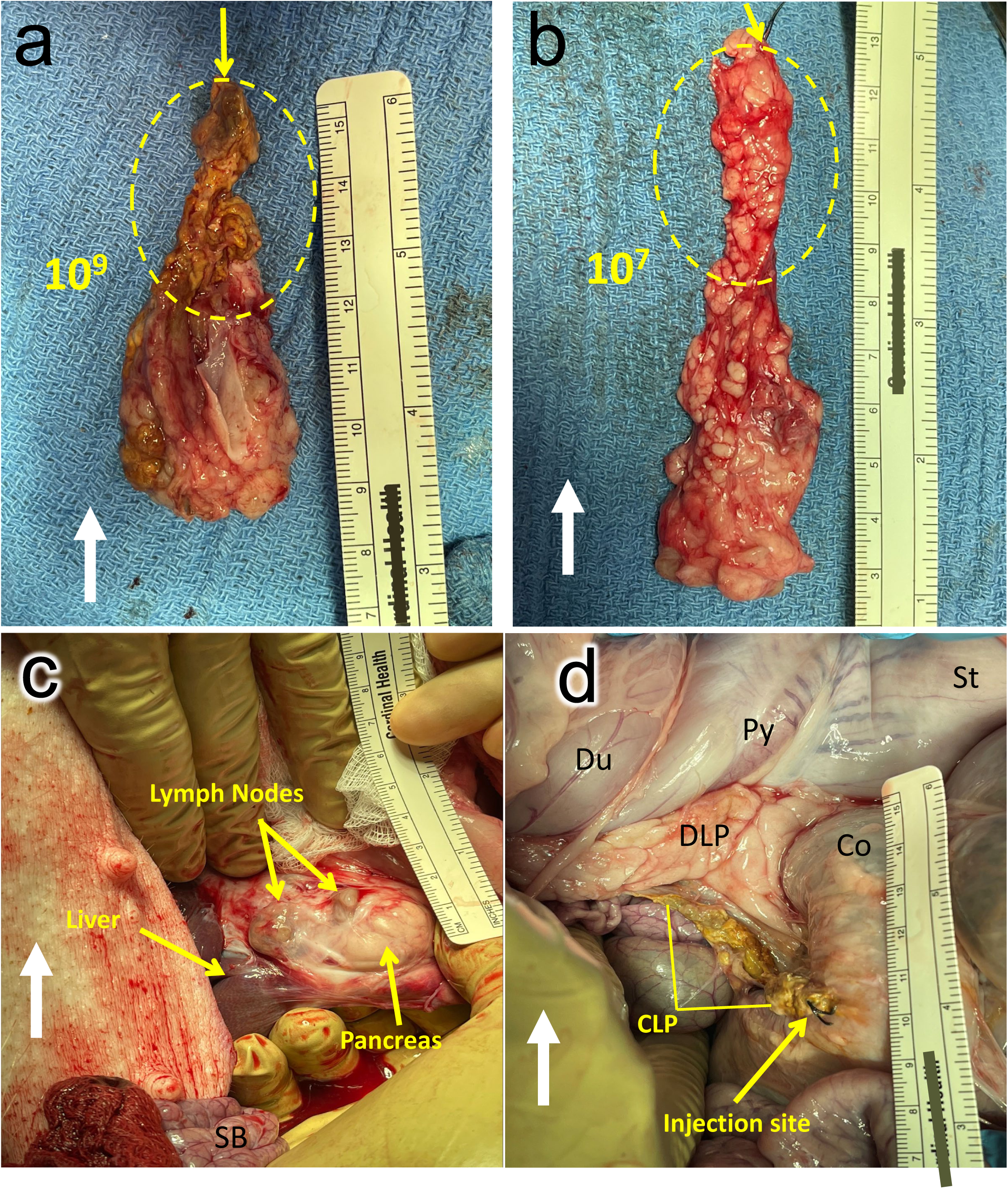
*In vivo* injection of Ad-K8-Cre into the duct of the connecting lobe of the pancreas in the Oncopig. (a) Identification of the Connecting Lobe of the Pancreas (CLP): Image shows the exposure of the CLP in the Oncopig. The stomach (St), pylorus (Py), and duodenum (Du) are retracted to enhance visibility of the surgical site. The surgeon’s finger (asterisk) isolates the CLP from the adjacent duodenal lobe of the pancreas (DLP). Du = duodenum; Co = colon; SB = small bowel; white arrow indicates cephalad orientation. (b) Continued Dissection of the Pancreas: Following further dissection, the junction between the CLP and DLP is isolated. A looped silk suture (asterisk) is placed around the junction for retraction. (c) Division and Preparation for Injection: The CLP is separated from the DLP, with the proximal end ligated using a silk suture (asterisk). The distal end of the CLP is prepared with an exposed duct, subsequently cannulated using a 22-gauge catheter. (d) Injection of Ad-K8-Cre Vector: The Ad-K8-Cre vector is injected into the duct of the CLP using a syringe, targeting the pancreatic ductal epithelium to induce Cre recombinase expression under the control of the Keratin 8 promoter.

After 2 months post-injection, all Oncopigs survived (Table 1). Oncopigs injected with 10^9^ to 10^10^ pfu of vector exhibited signs of pancreatic atrophy in the connecting lobe (i.e., the injection region) with small (<5 mm) clustered lumps (nodularity) and with enlarged (>1 cm), firm lymph nodes around the pancreas (Fig. 2a, c, d). On the other hand, Oncopigs injected with vector doses ranging from 10^7^ to 10^8^ pfu showed conserved gross morphology of the pancreas (Fig. 2b). At macroscopic level, the injection region in Oncopigs treated with 10^7^ to 10^8^ pfu showed neither tangible damage nor change. While there were lymph nodes present around the pancreas as well, there was an absence of discernible masses or marked discoloration to suggest atypical growth or inflammation with the lower dose Oncopigs (Fig. 2b, c). All subjects gained weight appropriately during the eight-week observation, and there were no differences in weight at any time point among the four dosage groups (Table 1). Blood testing (hemoglobin, hematocrit, amylase, lipase, carcinoembryonic antigen, CA 19-9, and alpha-fetoprotein) prior to induction vs. prior to necropsy revealed no differences in any group (Table 2).

**Table 1.** Oncopig weights.

| Ad-K8-Cre dose (pfu) | Survival post-induction | Mean weight (Kg) |  |  |  |  |
| --- | --- | --- | --- | --- | --- | --- |
|  |  | T<sub>0</sub> | 2 wk | 4 wk | 6 wk | Necropsy (8 wk) |
| 10<sup>7</sup> | 8 wk | 27.5 | 26.8 | 34.5 | 38.8 | 45.8 |
| 10<sup>8</sup> | 8 wk | 25.5 | 25.3 | 28.3 | 35.5 | 40.3 |
| 10<sup>9</sup> | 8 wk | 20.0 | 25.8 | 27.5 | 27.0 | 28.5 |
| 10<sup>10</sup> | 8 wk | 24.3 | 28.8 | 34.3 | 38.0 | 48.1 |
| *\*p value* |  | 0.5 | 0.8 | 0.6 | 0.6 | 0.3 |

\*ANOVA across all four dose groups at the given time point.

**Table 2.** Oncopig serum testing.

| Adcre | Sex | HG |  | HCT |  | AMYLASE |  | LIPASE |  | CEA |  | CA 19-9 |  | AFP |  |
| --- | --- | --- | --- | --- | --- | --- | --- | --- | --- | --- | --- | --- | --- | --- | --- |
|  |  | T0 | Necropsy | T0 | Necropsy | T0 | Necropsy | T0 | Necropsy | T0 | Necropsy | T0 | Necropsy | T0 | Necropsy |
| 10^9 | M | NA | 8.5 | NA | 30 | NA | 669 | NA | 3 | NA | 0.1 | NA | 1 | NA | 0.5 |
|  | F | 9.7 | 9.2 | 34.9 | 33 | 1520 | 1388 | 19 | 18 | 0.1 | 0.1 | 1 | 1 | 0.5 | 0.5 |
| 10^8 | M | 11 | 11.1 | 39.3 | 38.6 | 1015 | 1095 | 4 | 3 | 0.1 | 0.1 | 1 | 1 | 0.5 | 2.5 |
|  | F | 11 | 9.4 | 40.3 | 33.6 | 1452 | 1749 | 27 | 3 | 0.1 | 0.1 | 1 | 1.4 | 0.8 | 1 |
| 10^7 | M | 9.3 | 10.6 | 32.9 | 37.3 | 573 | 739 | 8 | 21 | 0.1 | 0.1 | 2 | 1 | 0.5 | 1 |
|  | F | 10.3 | 10.4 | 36.4 | 36.9 | 1551 | 1865 | 4 | 6 | 0.1 | 0.5 | 1 | 1 | 0.5 | 0.5 |
| 10^10 | M | 9.1 | clotted | 34.2 | clotted | 674 | 799 | 3 | 3 | 0.1 | 0.3 | 1 | 1 | 0.5 | 0.5 |
|  | F | 10.4 | 10.2 | 38.2 | 36.9 | 1477 | 2556 | 3 | 3 | 0.1 | 0.1 | 1 | 1 | 0.5 | 0.5 |

Hg = hemoglobin; Hct = hematocrit; CEA = carcinoembryonic antigen; CA 19-9 = cancer antigen 19-9; AFP = alpha fetoprotein; T0 = immediately prior to induction.

### Microscopic Morphology

Two months after Ad-K8-Cre delivery, histological analysis of pancreas from the proximal connecting lobe, i.e., the region of injection (Fig. 3), showed architectural disruption and cellular atypia for all vector doses tested, whereas pancreatic samples remote from the injection site (i.e., located more than 10 cm away from the connecting lobe) exhibited normal pancreatic architecture. In this study, each Oncopig served as its own control. That is, the pancreas in these swine was large enough so that a biopsy of normal appearing pancreas more than 10 cm away from the vector injection region (i.e., the proximal connecting lobe) was possible in each subject. So, samples of treated *versus* untreated pancreas could be acquired from the same pig. The above histologic changes were accentuated in the 10^9^ and 10^10^ vector dose (Fig. 3a).

**Figure 3:**
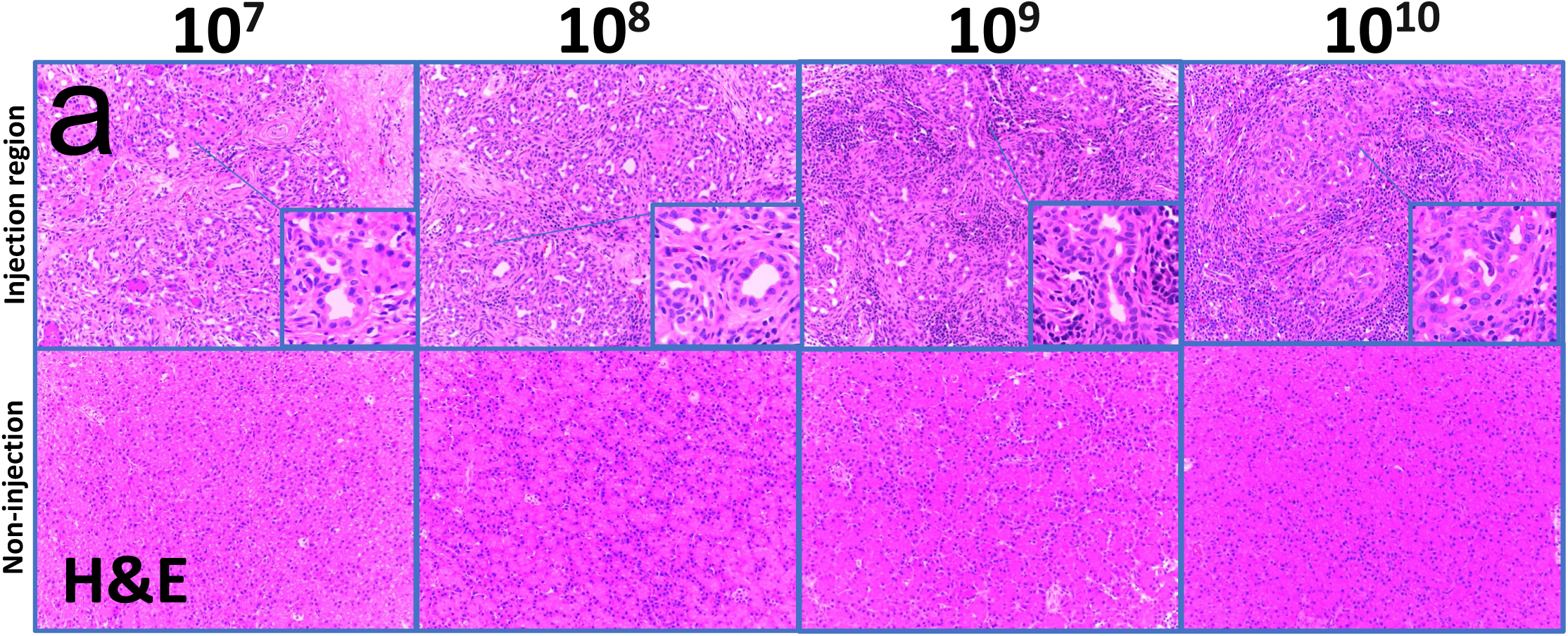
Histological and Immunohistochemical Analysis of Pancreatic Tissues Post Ad-K8-Cre Administration. Hematoxylin and Eosin (H&E) Staining of Pancreatic Tissue: Sequential panels display H&E-stained sections of pancreatic tissue from Oncopigs injected with increasing doses of Ad-K8-Cre (10^7^ to 10^10^ plaque-forming units, pfu). The top row shows histological images from the injection site regions, revealing dose-dependent alterations in tissue architecture and cellular morphology. Insets provide higher magnification views of cellular details. The bottom row displays tissues from regions remote (>10 cm) from the injection site, illustrating preserved normal pancreatic architecture and serving as internal controls for the localized effects of Ad-K8-Cre administration.

### Proliferation

Ductal structures within the injection region were populated by KI67 and PCNA-positive cells, which was evident at all Ad-K8-Cre concentrations tested (Fig. 4a-d). While the 10^7^ to 10^9^ pfu concentrations displayed a ductal proliferation pattern, the 10^10^ pfu showed a loss of distinct circular or ovoid proliferation areas, with stronger, more dispersed, and homogenous staining pattern throughout the sample. Pancreas remote from the injection region maintained normal microscopic morphology, including uniform tissue structures and limited KI67 staining (Fig. 4a-b).

**Figure 4:**
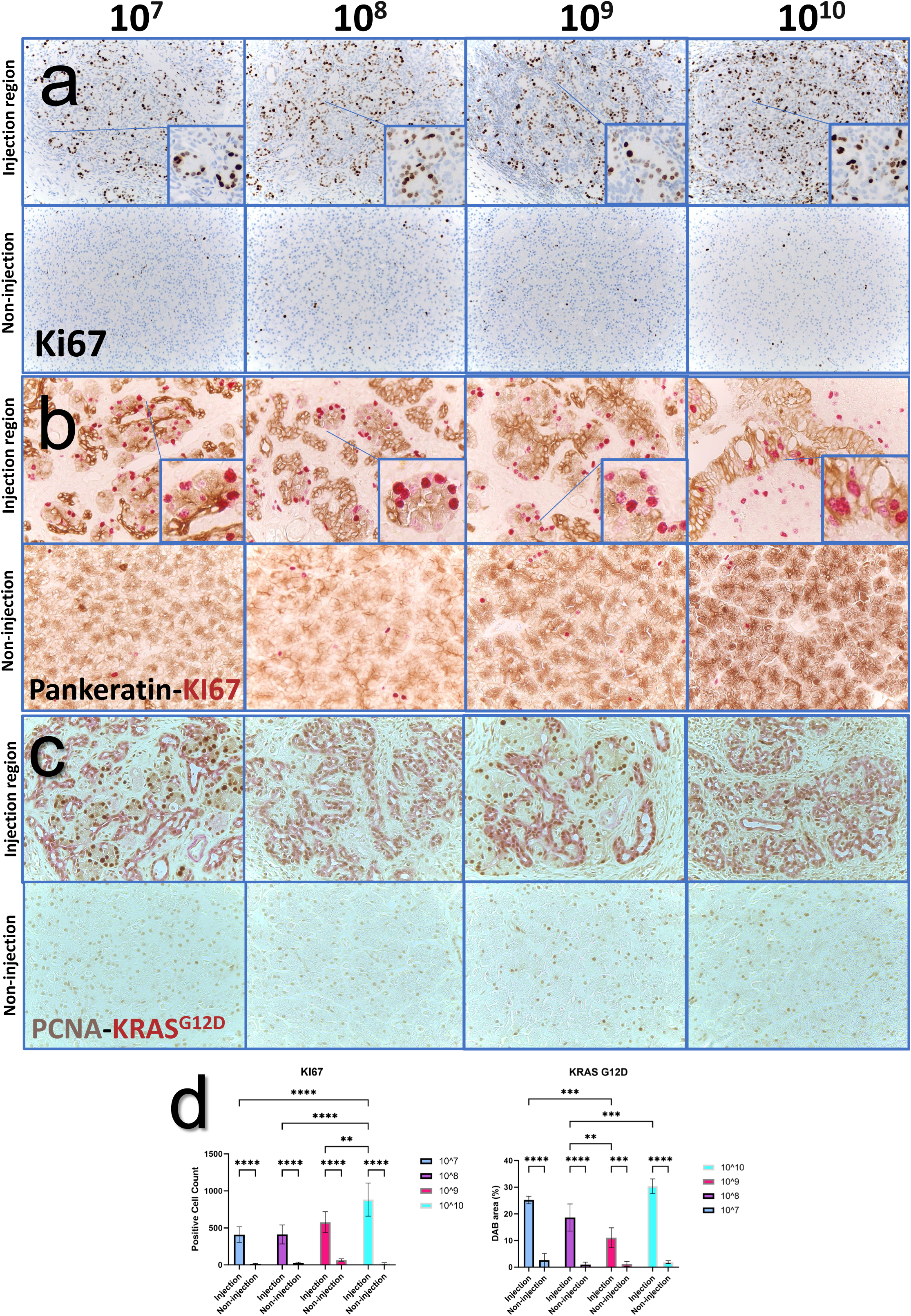
Immunohistochemical Analysis of Proliferation and Oncogenic KRAS Expression in Pancreatic Tissues Post Ad-K8-Cre Injection. (a) KI67 Staining: Representative images of KI67 immunostaining in pancreatic sections from Oncopigs injected with varying doses of Ad-K8-Cre (10^7^ to 10^10^ pfu). The top row shows the injection site regions with increased KI67-positive cells, indicating proliferation. Insets provide magnified views. The bottom row shows non-injection site regions, maintaining minimal KI67 expression as internal controls. (b) Double Staining for Pankeratin and KI67: Dual immunohistochemistry for Pankeratin (brown) and KI67 (magenta) at the injection sites in the pancreas highlights ductal epithelial cell proliferation. Insets display close-up views of proliferating ductal cells. Non-injection sites in the bottom row exhibit intact ductal architecture without significant KI67 labeling. (c) Double Staining for PCNA and KRAS^G12D^: PCNA (brown) and mutant KRAS^G12D^ (red) staining in pancreatic sections post-injection. The injection sites show positive staining for KRAS^G12D^ within proliferative ductal epithelial cells, confirming successful recombination and oncogenic KRAS expression. The bottom row includes non-injection sites as negative controls, showing no KRAS^G12D^ expression. (d) Quantification of KI67 and KRAS^G12D^ Expression: Bar graphs quantify KI67-positive cell counts (left) and KRAS^G12D^-positive DAB area (right) across different doses at both injection and non-injection sites. Statistical significance is indicated: ****p<0.0001, ***p<0.001, **p<0.01, *p<0.05.

In addition, the following histologic characteristics were observed within the injection region:

i. *Acinar Cell Loss and Ductal Proliferation.* While the Ad-K8-Cre specifically drove expression in ductal cells, we observed loss of acinar morphology within the injection region, along with a significant increase in ductal cell proliferation (Fig.3a, 4a-c, Supplementary S3a-c).
ii. *Intestinal metaplasia in the large ducts*. In some regions of the larger pancreatic ducts, the flat ductal epithelium showed the appearance of goblet cells (Supplementary S4a).
iii. *Inflammatory response*. At the two-month time point, the inflammatory response was assessed by identifying CD45-positive leukocytes, CD3-positive T lymphocytes, and MPO-positive neutrophils. We identified increased T lymphocytes with lower neutrophils infiltration (Fig. 5a-d).
iv. *Healing vs. regeneration vs. neoplasia*. Based on the H&E morphology and the immune cell staining, it was not possible to determine whether the above histologic changes were representative of a response to injury that would generate a scar (i.e., wound healing), or if a regenerative response would ensue, or if there was microscopic neoplasia present which would produce a tumor.

**Figure 5:**
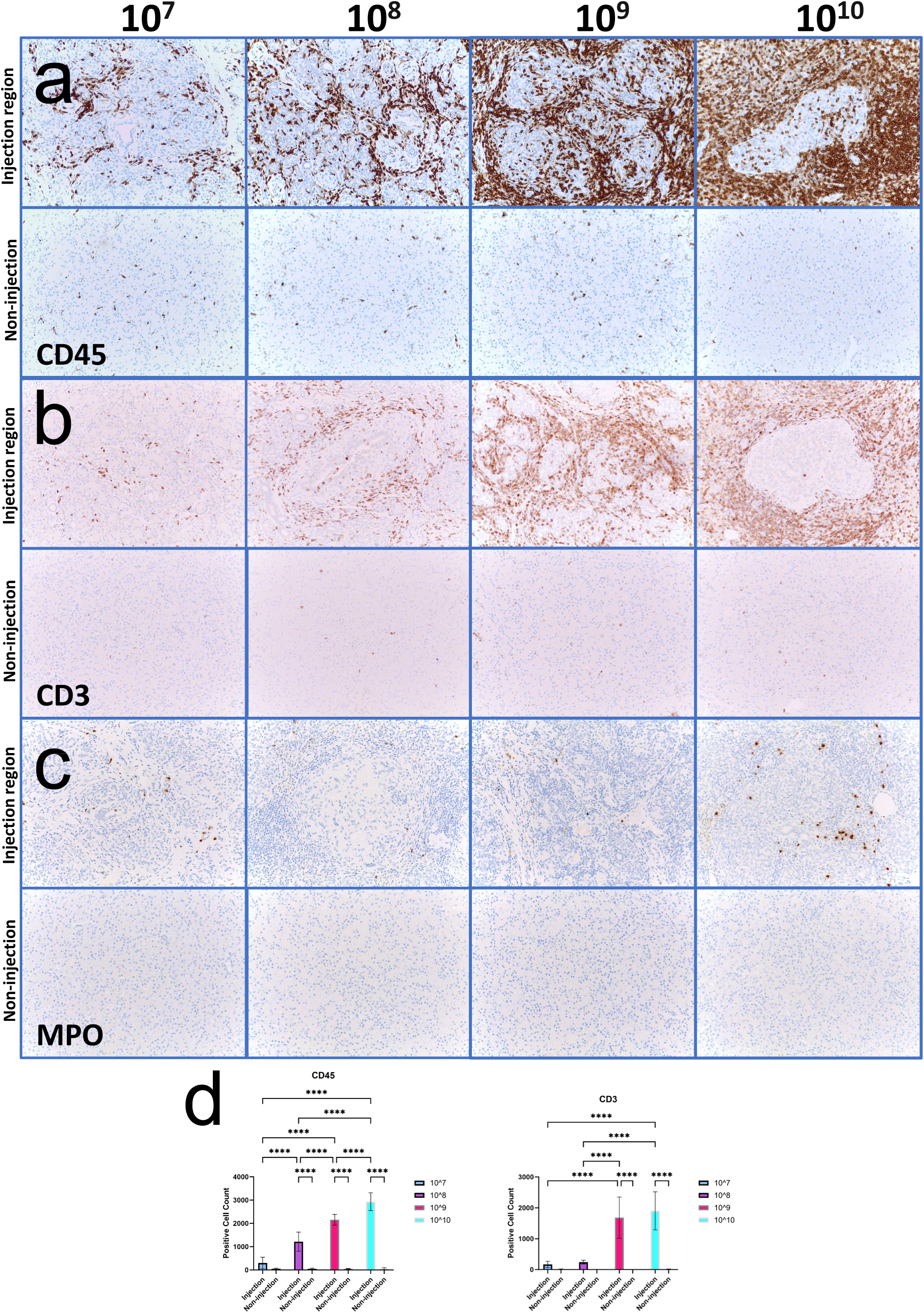
Immune Cell Infiltration in Pancreatic Tissues Following Ad-K8-Cre Injection. (a) CD45 Immunostaining: Representative images of CD45 immunohistochemistry on pancreatic sections from Oncopigs injected with increasing doses of Ad-K8-Cre (10^7^ to 10^10^ pfu). The top row displays injection site regions, where CD45-positive immune cell infiltration is dose-dependent. The bottom row shows non-injection sites, with minimal to no CD45-positive cells as internal controls. (b) CD3 Immunostaining: Immunostaining for CD3-positive T cells in pancreatic tissue at injection sites across different doses. Increased T cell presence is observed with higher Ad-K8-Cre doses, particularly in peri-ductal regions. Non-injection sites (bottom row) display minimal CD3 staining. (c) MPO Immunostaining: MPO staining illustrates neutrophil infiltration at injection sites, showing a sparse but dose-dependent increase in MPO-positive cells at higher Ad-K8-Cre doses. Non-injection regions exhibit no significant MPO staining. (d) Quantification of Immune Cell Markers: Bar graphs quantify CD45-positive and CD3-positive cell counts across different doses in injection and non-injection regions. Statistical significance is indicated as follows:****p<0.0001, ***p<0.001, **p<0.01, *p<0.05.

### Targeted KRAS^G12D^ Expression Within Epithelium

After 2 months post Ad-K8-Cre injection, we observed a targeted induction of the KRAS^G12D^ expression within the epithelium (primarily in the ductal cells) within the injection region (Fig. 4c). These staining patterns were particularly evident in the areas with disrupted architecture and complex ductal proliferation. This staining confirms that the Ad-K8-Cre delivery induced KRAS^G12D^ expression within the epithelial cells. Interestingly, regions that tested positive for the KRAS^G12D^ mutation also displayed a corresponding rise in KI-67 expression. We confirmed that proliferation is associated with cells expressing KRAS^G12D^ by performing double staining for PCNA and KRASG12D, as shown in Figure 4c. This underscores the relative quiescence of healthy pancreas compared to the aggressive growth pattern observed in the injected region.

### Cytokeratin expression In Transformed, Proliferative Regions Suggest Epithelial Origin

We utilized epithelial markers, (Pan-keratin, CAM5.2 and Cytokeratin 7) to confirm the epithelial nature of the transformed pancreatic cells. Pan-keratin (Fig. 4b), CAM5.2 (Supplementary S3b), and Cytokeratin 7 (Supplementary S3c) stained strongly in cells corresponding to those transformed by KRAS within the injection region (Supplementary S3a). We observed the highest staining intensity in samples from Oncopigs injected within 10^9^ and 10^10^ concentrations, highlighting the effectiveness at elevated doses and suggesting a dose-response relationship. The epithelial markers were reduced in pancreas remote from the injection, emphasizing the targeted expression of Cre recombinase, and suggesting that the KRAS^G12D^-expressing cells maintained epithelial lineage post-transformation.

### Desmoplasia, Matrix Deposition, and Mucin Production

Histologic features of stromal reaction to Ad-K8-Cre transfection were characterized with vimentin, α-SMA, and Masson’s trichrome staining (Fig. 6a-c). Increased collagen deposition consistent with desmoplasia was found within the injection region for all concentrations (Fig. 6a). In particular, there was a substantial fibrotic and desmoplastic stromal reaction at the 10^9^ and 10^10^ concentrations (blue staining in Fig. 6a). α-SMA (smooth muscle actin) immunohistochemistry demonstrated a vector dose-dependent increase in staining (Fig. 6b), consistent with a stromal reaction within the injection region. Conversely, α-SMA staining was minimal within pancreas remote from the injection region (Fig. 6b). In addition, there was increased vimentin staining within the stroma surrounding proliferative ductal cells within the injection region, also consistent with desmoplasia (Fig. 6c). The 10^9^ to 10^10^ vector doses had the most intense and widespread vimentin staining; pancreas remote from the injection region had minimal vimentin signal. There also was an increase in acidic mucin staining in the ductal epithelium and stroma within the injection region (purple-colored staining in Fig. 7), with an apparent dose-dependency on vector dose. In contrast to the injection region, remote healthy pancreas displayed minimal staining.

**Figure 6:**
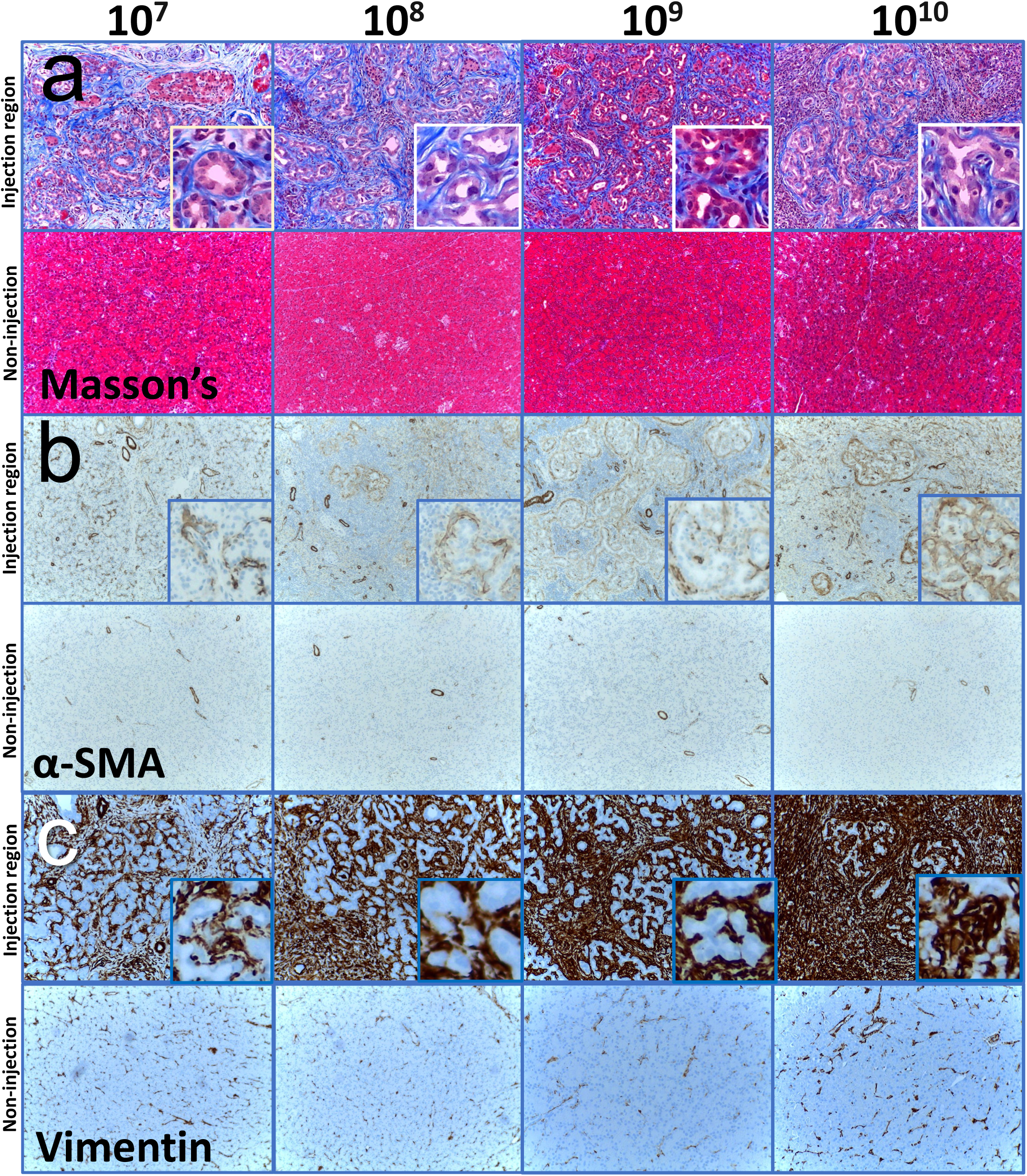
Evaluation of Stromal Reaction in Pancreatic Tissues Post Ad-K8-Cre Injection. (a) Masson’s trichrome staining of pancreatic tissues. Top row: Increased blue staining with the injection regions across increasing vector doses (10^7^ to 10^10^ pfu). Bottom row: Minimal staining in pancreas remote from the injection regions, showing normal collagen levels. (b) α-SMA immunohistochemistry used to detect stromal cells involved in the fibrotic response. Top row: Enhanced α-SMA staining within the injection regions with increasing vector doses. Bottom row: staining within non-injection regions. (c) Vimentin immunostaining. Top row: Marked increase in vimentin staining within the injection regions, particularly notable at doses 10^9^ and 10^10^ pfu. Bottom row: Sparse vimentin staining from the non-injection regions.

**Figure 7:**
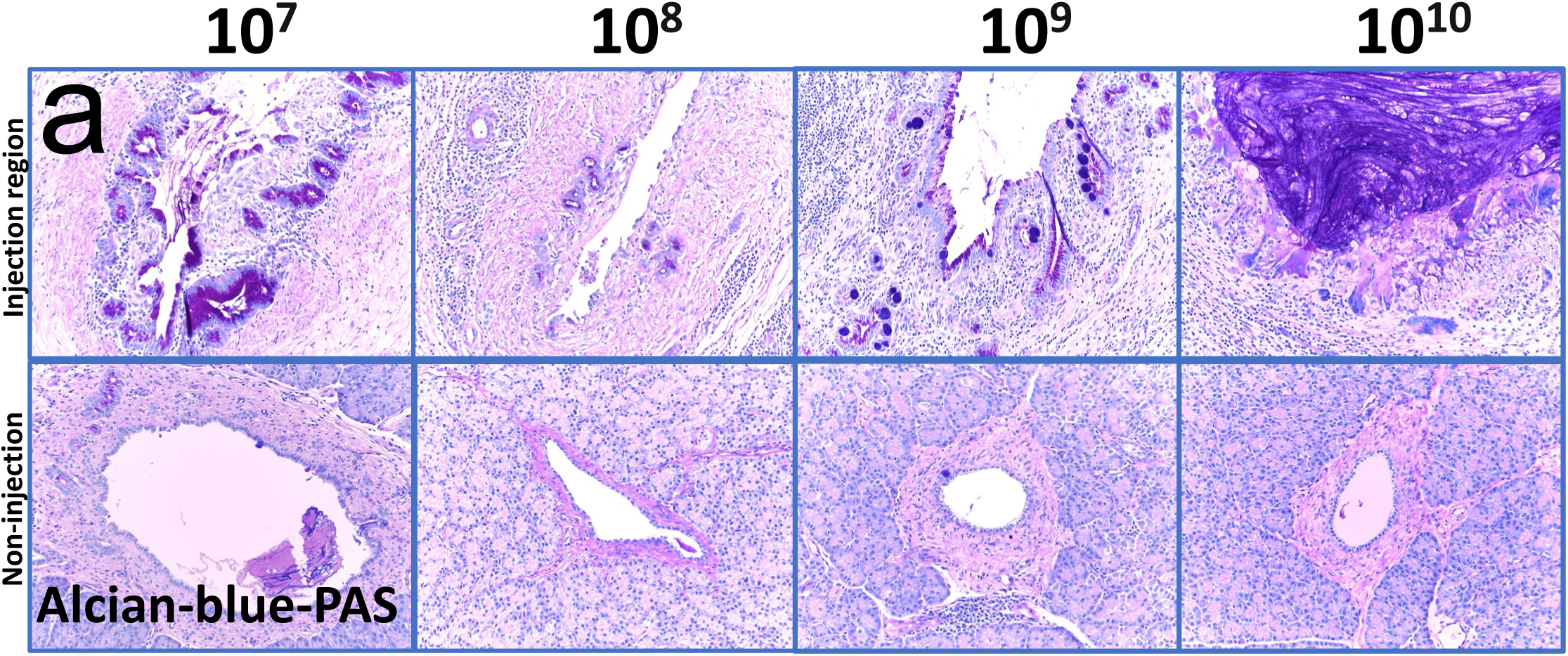
Alcian Blue-PAS Staining of Pancreatic Tissues Post Ad-K8-Cre Injection. Top Row: progression of acidic mucin production with the injection regions of Oncopigs administered with increasing doses of Ad-K8-Cre (10^7^ to 10^10^ pfu). Non-injection Region (Bottom Row): limited Alcian Blue-PAS staining.

### Comparative Human Microscopy

Examination of four human pancreatic cancer samples (Supplementary Fig. S5) demonstrated considerable architectural disruption, indicative of advanced disease. Human pancreatic cancer tissue also was characterized by a desmoplastic reaction, with increased collagen and vimentin deposition. These changes were comparable to the those observed with vector induction in the Oncopig model, particularly at the higher vector doses.

## DISCUSSION

### Intraductal Injection for Vector Delivery

The unique, tri-lobed anatomy of the porcine pancreas allows for isolation and injection of a large pancreatic duct (within the connecting lobe) without major physiologic impact to the subject (6, 17). This anatomic arrangement, along with the large subject size, allows for unique opportunities to study pancreatic neoplasia in swine. For example, the swine’s unique anatomy allows for precise delivery of the adenoviral vector, Ad-K8-Cre. Intraductal administration of the vector ensures that the adenovirus will bathe the pancreatic epithelium, maximizing the potential for targeted gene activation. These theoretical advantages were borne out by the results of this study. For example, the injection region was the open end of the primary duct to the connecting lobe where the cannula was inserted (Supplementary Fig. S2) and we presume it diffused into the regional pancreas (i.e., proximal connecting lobe) via the branching ductal system. Endoscopic Retrograde Cholangiopancreatography (ERCP) offers advantages for pancreatic intraductal injections compared to traditional surgical methods. Its minimally invasive nature reduces the need for large incisions, leading to shorter recovery times and fewer postoperative complications (18-21). Additionally, ERCP offers real-time imaging allowing precise targeting of the pancreatic ducts (22, 23). The endoscopic approach is less invasive, but often presents challenges in terms of investigator accessibility (i.e., equipment and operator expertise requirements). It is not clear whether the duct to the connecting lobe can be always reliably cannulated via ERCP. With the open surgical approach, our success rate in cannulation of the duct to the connecting lobe was 100%. Our laparotomy-based approach, involving dissection and direct visualization of the pancreatic duct, requires no special equipment other than what typically can be found in the typical Comparative Medicine Department, and can be taught to individuals who has veterinary operative experience.

### Dose Dependence

At a lower vector concentration (10^7^ pfu of virus per injection), Ad-K8-Cre promoted epithelial proliferation while preserving ductal structural integrity. This suggests that the adenoviral vector can precisely target the epithelial cells without disrupting the ductal architecture with only a mild desmoplastic response. In contrast, the highest vector concentration tested (10^10^) produced a marked increase in disorganized epithelial proliferation (KI67 staining), and a robust desmoplastic reaction characterized by extensive immune infiltration, collagen deposition and α-SMA expression. This suggests that at elevated concentrations, Ad-K8-Cre not only impacted epithelial cells but also triggered a desmoplastic/fibrotic response that could potentially modify the tumor microenvironment. This dose-dependent response ultimately may prove useful for diverse research efforts on pancreatic cancer.

### Intestinal metaplasia and Goblet Cells

Goblet cells are specialized epithelial cells found primarily in the mucosal lining of the intestines and respiratory tract, where they play a crucial role in secreting mucus to protect and lubricate the mucous membranes (24). Intestinal metaplasia occurs when epithelial cells in one area of the body transform into intestinal-type epithelial cells, typically as a response to chronic irritation or inflammation (25). The prototypical example of intestinal metaplasia is Barrett’s esophagus, in which the stratified squamous epithelium of the esophagus transdifferentiates into columnar-type intestinal epithelium, which is associated with an increased risk for esophageal carcinoma (26). Pancreatic ducts do not naturally contain goblet cells; their appearance in our study is consistent with intestinal metaplasia. Similar to acinar-to-ductal metaplasia this type of metaplasia can be triggered by chronic inflammation or injury (25, 27). However, as in Barrett’s esophagus, the appearance of goblet cells in the pancreatic ducts could signify an increased risk for epithelial neoplasia.

### Large animal modeling of pancreatic cancer

Over the past 30 years, murine have helped to improve the understanding of gene function, cellular transformation, and other aspects of pancreatic cancer biology. One of the most extensively studied murine model is the KPC model (Kras^G12D^/+; Trp53^R172H^/+; Pdx1-Cre)(12). This model incorporates both KRAS^G12D^ and p53^R172H^ mutations driven by Pdx1-Cre, targeting pancreatic progenitor cells and producing tumors that mimic the aggressive nature and metastatic behavior of human PC model. However, murine models, including the KPC, represent limitations when scaling to applications that require a human-sized subject, such as testing neoadjuvant therapies or surgical resections. On the other hand, the targeted expression of the KRAS^G12D^ mutant to pancreatic ductal epithelium within the Oncopig model underscores the potential of large animal models to advance PDAC research. Porcine pancreatic cancer models could provide a realistic and clinically relevant platform for preclinical assessment of novel therapeutics and diagnostics, and potentially may permit mechanistic studies of early tumor progression (6, 9). In contrast to a recent investigation on a porcine model of pancreatic cancer (6) which used a nonspecific adenoviral vector (Ad5CMVCre-eGFP,) in the Oncopig and reported fulminant tumor growth, short survival times, and mixed tumor pathology, the present study on the use of the Ad-K8-Cre induction vector demonstrated enhanced specificity for pancreatic epithelial cells, which likely produced the 100% 2-month survival. The implication of this result is that neoplasia induced by the K8-driven vector will be a slower growing tumor model which could present opportunities for early diagnostic and interventional technologies, along with mechanistic studies of tumor progression.

In addition, the genetic and physiological similarities between pigs and humans could better enable the study of tumor progression and therapeutic responses in a setting that is far more comparable to human biology than murine models. These porcine models allow researchers to explore complex multimodal treatments, including surgery, chemotherapy, and localized therapies, which are more difficult to evaluate in smaller animals like mice (28). Furthermore, the size of the pig allows for the use of human-sized medical equipment, facilitating translational research in diagnostic and therapeutic device development (4, 9). Unlike murine models, which often fail to replicate the full scope of human immune responses and tumor microenvironments, porcine models provide a robust platform to study interactions between cancer cells and the immune system, a critical component in the development of immunotherapies (4, 28).

The question of whether true neoplasia or carcinoma was produced in this study is still open. Careful and independent pathologic analysis was inconclusive on this question. There was obvious evidence of mutant KRAS expression along with increased proliferation and desmoplasia typical of PDAC. Perhaps an early *in situ* transformative process was being observed; however, we cannot be conclusive with the present data. A longer observation period than the two months used in the present report may be needed to resolve this question.

Another open question from this study is the role of the host immune response to injection of the vector into the pancreas. It was demonstrated in previous work (6) that the vector itself has minimal pathologic effect; it was only when the vector expressed Cre recombinase (driven by the CMV promoter) that there was strong evidence of transformation accompanied by an overwhelming immune response. The immune response to epithelial-restricted Cre recombinase expression in this report was qualitatively less than in the prior study of unrestricted Cre recombinase expression (6) but a T-cell response was still present in the current report. Perhaps the Oncopig’s T-cell response to nascent ductal transformation will provide an opportunity for mechanistic studies on host immune response to PDAC.

### Limitations

A formal sample size calculation was not conducted, as this was an exploratory study. We acquired as many Oncopigs as our budget allowed to maximize the robustness of our observations. This approach provided sufficient data for initial analysis and hypothesis generation for future studies with larger, statistically powered sample sizes.

Each Oncopig served as its own control. The pancreas of each animal was divided into two regions: the injection (experimental region) and a non-injection region (control region), located more than 10 cm away from the injection region within the same pancreas. This setup allows for direct within-subject comparison, minimizing inter-animal variability and strengthening the reliability of observed effects. Groups and controls were evaluated under identical conditions to ensure consistency across experimental outcomes.

## CONCLUSION

Our study demonstrated that intraductal administration of the adenoviral vector Ad-K8-Cre had a qualitative dose-dependent effect on epithelial cell proliferation and stromal reaction within the injection region; the latter effect was reminiscent of human PDAC. These histologic findings were related to expression of the KRAS^G12D^ mutation. While we have no incontrovertible evidence for neoplastic transformation in the present study, we believe that future work will demonstrate that the histologic events noted in the present study were indicative of early stage PDAC.

## DECLARATION

The datasets used and/or analyzed during the current study available from the corresponding author on reasonable request.

## MATERIALS AND METHODS

### Animal Welfare

The animals utilized for this report were maintained and treated in accordance with the *Guide for the Care and Use of Laboratory Animals* (8^th^ ed.) and in accordance with the Animal Welfare Act of the United States (U.S. Code 7, Sections 2131 – 2159). The animal protocol pertaining to this manuscript was approved by the Institutional Animal Care and Use Committee (IACUC) of the University of Nebraska Medical Center (ID number 19-053-FC). All procedures were performed in animal facilities approved by the Association for Assessment and Accreditation of Laboratory Animal Care International (AAALAC; www.aaalac.org) and by the Office of Laboratory Animal Welfare of the Public Health Service (grants.nih.gov/grants/olaw/olaw.htm). All surgical procedures were performed under isoflurane anesthesia, and all efforts were made to minimize suffering. Euthanasia was performed in accordance with the AVMA Guidelines (American Veterinary Medical Association, 2020. AVMA guidelines for the euthanasia of animals: 2020 edition. American Veterinary Medical Association). Design and conduct of animal experiments was guided by the Animal Research: Reporting of In Vivo Experiments (ARRIVE) standards (Supplementary 6).

### Porcine Subjects

Transgenic Oncopigs (Oncopig Model, or OCM; LSL-*KRAS*^G12D^-IRES-*TP53*^R167H^, Supplementary S2c). were purchased from Sus Clinicals (susclinicals.com). The OCM subjects were a hybrid of Minnesota minipigs and domestic pigs. The genotype of each porcine subject was confirmed with PCR upon subject delivery. Swine were housed two littermates per pen, except for one week of individual housing post-laparotomy (but with contact through the pen grating), in order to prevent wound cannibalism. Swine were fed ad lib with standard hog feed (Purina Nature’s Match® Sow and Pig Complete Feed; www.purinamills.com). The basic experimental design included a ≥1 week acclimatization period after subject delivery to the research facility. Each subject then underwent an induction procedure (laparotomy under general anesthesia; one major survival procedure), followed by observation for 2 months.

### Overview

The technique of vector administration into the Oncopig pancreas is based on a previous description (6). Briefly, our laparotomy procedure initiated with a ventral midline incision, starting just inferior to the xiphoid process, and extending to the umbilicus in females and approaching the urethral meatus in males. After retracting the small and large bowel caudally, exposing the duodenal segment of the pancreas. The connecting lobe of the pancreas was transected just below its intersection with the duodenal lobe. Using surgical loupes (3.5x magnification), the duct within the connecting lobe was cannulated using a 22g Angiocath. The specified concentration of Ad-K8-Cre (from 10^7^ to 10^10^ pfu) in 100-150 µL was injected into this duct, which was immediately ligated with 3-0 silk after the injection in order to prevent back flow of the injectate out of the duct. The pancreas remote (>10 cm) from the injection site was used as a same-subject control, providing a baseline for comparison with the treated regions.

### Operative Details

All Oncopigs underwent tumor induction in the same fashion. The Oncopigs were pre-anesthetized using an intramuscular injection of Telazol®-Ketamine-Xylazine (2/4/2 mg/kg, respectively) placed supine on the procedure table, and then intubated with 6-7 French endotracheal tube. Anesthesia was maintained with 1% Isoflurane inhalation. Vitals were monitored noninvasively and maintenance fluid, (Lactated ringers’ solution) was given throughout the case (200-300 mL total).

The skin was shaved and prepped with chlorohexidine in alcohol, weight-based bupivacaine was injected subcutaneously around the planned upper abdominal midline, and abdomen was draped in sterile fashion. An incision was made below the xiphoid extending down past umbilicus in females and proximal to the urethral meatus in males with #10 blade. This was then carried down through subcutaneous fat, facia and muscle to the pre-peritoneal space using electrocautery. The peritoneum was carefully incised to enter the greater sac and Bookwalter retractor was assembled and used to provide adequate abdominal exposure.

The small and large bowel were retracted caudally, and the stomach was grasped at the pylorus for reflection antero-superiorly to expose the duodenal lobe of the pancreas. Pancreatic adhesions to the small/large bowels were cut and a window between the connecting lobe of the pancreas and pancreatic head (duodenal lobe) was created. After the proximal connecting lobe of the pancreas had been dissected circumferentially, the proximal end of the connecting lobe of pancreas was tied off with 3-0 silk flush with the duodenal lobe.

The connecting lobe then was transected 0.5-1 cm distally from the silk ligature. With the aid of surgical loupe magnification (3.5x) the pancreatic duct in the distal connecting lobe was identified and cannulated with 22 g Angiocath. The injectate with the desired pfu of Ad-K8-Cre was injected into the pancreatic duct, the catheter was withdrawn, and the distal connecting lobe of pancreas was immediately ligated with 3-0 silk suture.

The peritoneum was reapproximated with 3-0 Vicryl suture; the midline abdominal fascia was closed with 0-Maxon suture (5 mm tissue bite, 2-3 mm stitch interval); the deep dermal layer was closed with 4-0 Vicryl suture; and a subcuticular closure with 4-0 Vicryl suture on a cutting needle. Dermabond adhesive glue was then placed over the incision. The Oncopigs were extubated after confirmation of spontaneous breathing and continued to be monitored until oxygen saturations remained greater than > 90 on room air.

### Postoperative Care and Observation Period

Oncopigs were housed in individual pens until post-op day 4 to prevent wound cannibalism, after which they were regrouped with their respective sex. All animals were monitored carefully for 3 days and were kept NPO until postoperative day 2 once, at which time feeding was reinitiated if the subject was clinically stable. The swine continued to be assessed daily for signs of deterioration until term. Criteria for early removal from the study and euthanasia were symptoms of failure to thrive (anorexia, lethargy, decreased movement, abnormal breathing, or other signs of distress) or sepsis (fever, wound disruption, or drainage)

### Euthanasia and Necropsy

At 2 months post AdCre induction, the Oncopigs underwent necropsy. Anesthesia and access to the abdomen was obtained in a fashion similar to the above, taking care to lyse adhesions and avoid bowel injury. At the time of euthanasia, inhalational isoflurane was increased to 5% via the ventilator. The prior midline incision was reopened. Inferiorly this incision was extended in paramedian fashion to avoid midline structures (such as the urethra in males). The completed necropsy incision extended from xiphoid process to the pelvic inlet. The bilateral thoracic cavity then was entered by transversely incising the diaphragm just inferior to the xiphoid process. The intrathoracic portion of the inferior vena cava was easily identified as it emerged from the liver in the posterior mediastinum. Phlebotomy was performed from the cava, and then Fatal-Plus® (pentobarbital sodium, 390 mg/mL; 1 mL per 4.5 kg body weight) was administered by caval injection. Two minutes after administration of Fatal-Plus®, the inferior vena cava was transected just above the diaphragm to exsanguinate the subject. A gross necropsy involving the lungs, heart, liver, kidneys, pancreas, intestines, bladder, and associated peritoneal surfaces then was performed. The pancreas and surrounding tissue then underwent gross evaluation and circumferential excision with tissue specimens collected from the injection region (connecting lobe), non-injection region (tail of pancreas), and lymph nodes for histopathologic analysis.

### Ex Vivo Pancreatic Duct Procedures

For the *Ex Vivo* tissue culture, pancreatic samples were processed as previously described with some modifications (29, 30). Briefly, porcine pancreases were harvested from wild type domestic pigs. The main pancreatic duct was carefully isolated from the harvested pancreas. The isolated pancreatic main duct was meticulously chopped into small pieces. The dissected pancreatic duct pieces were placed in a 6-well plate. Each well was filled with DMEM supplemented with 10% FBS (tetracycline-free). A dose of 10^9^ pfu of Ad-K8-Cre was added to each well. The plate was incubated overnight in a 5% CO2 incubator. Following incubation, the samples were fixed using 4% paraformaldehyde overnight. The fixed samples were then embedded in paraffin and sectioned at 5µm thickness. The sections were incubated overnight with an anti-recombinase antibody. Post incubation, the sections were stained using DAB to visualize the recombinase expression.

### Histology and Immunohistochemistry

For the histological analyses, pancreatic samples were processed and stained with hematoxylin and eosin (Hematoxylin Select Reserve Eosin Multichrome) as previously described (31). Briefly, Pancreatic tissues were harvested from porcine subjects. Two distinct samples were obtained from each subject: Injection region (Connective lobe where AdCre was administered) and non-injection region (pancreas distant from the injection region, particularly the tail or splenic lobe of the pancreas). Harvested samples were immediately fixed in cold 4% paraformaldehyde overnight. Post-fixation, the tissues were embedded in paraffin and serially sectioned at 5µm thickness. Three serial sections from both regions (injection and non-injection) per animal were prepared for IHC and histological analysis. We incubated sections with hematoxylin for 30 s, rinsed in distilled water, incubated for 30 s with eosin, rinsed again in distilled water, and dehydrated. The tissue sections were incubated overnight with the following primary antibodies: α-SMA (1:1000, Abcam ab5694), Anti-cytokeratin 8 (Invitrogen MA1-19037, 1:500), Anti-CAM5.2 (1:500, Roche REF#790-4555) Anti-KRAS G12D (Genetex GTX132407 1:200), Anti-KI67 (ab16667, 1:200), and Anti-vimentin (Roche REF#790-2917, 1:500). After incubation, the sections were treated with DAB to detect protein expression. For Masson’s Trichrome Staining (Masson Trichrome Stain Kit, #KTMTR EA), we incubated Weigert’s Iron Hematoxylin for 5 min, Biebrich Scarlet Acid Fuchsin solution for 15 min, differentiate in the phosphomolybdic acid solution for 15 min, Aniline Blue solution for 10 min and acetic acid solution for 5 min. Finally, we mounted the slices on Entellan®, digitally captured the images under bright field microscopy (32). For the mucins and polysaccharides analyses, pancreatic samples were processed and stained with Alcian Blue-Periodic Acid Schiff (PAS, Alcian Blue PAS Stain Kit, Liter #KTAPALT EA) as previously described (33). Briefly, Alcian Blue solution of pH 2,5 was made by dissolving Alcian Blue in distilled water and adding glacial acetic acid. The neutral red stain was made by dissolving 1 g of neutral red in 100 mL of distilled water and adding 0,1 ml of glacial acetic acid. In short, slides were deparaffinised and rehydrated, stained for 15 min in the Alcian Blue solution, rinsed for 5 min in tap water, counterstained with the Neutral Red solution for 1 min and then dehydrated, cleared, and mounted. All staining procedures were carried out in accordance with manufacturer’s recommendations.

### Pancreatic Transcriptomic analysis

For the single cell RNA sequencing analysis, publicly available healthy single cell samples (34, 35) were processed as previously described (36, 37) with some modifications. Briefly, cells with library size < 1000 UMI or few expressed genes were excluded from analysis and processed into the Seurat R package (v4.0). The Seurat objects and pipeline were used for downstream analyses and visualization: NormalizeData, ScaleData were used to calculate the comparable expression values; FindVariableFeatures were used to include the variable genes that contribute to the overall similarity/variability of cellular transcriptomic profiles; RunPCA, FindNeighbors, FindClusters, and RunUMAP were used to calculate the dimension-reduction coordinates for visualization and to perform unsupervised clustering. In the downstream analyses, we used Uniform Manifold Approximation and Projection (UMAP) coordinates to visualize the layout of the cells (34).

## SUPPLEMENTARY FIGURE LEGENDS

**Figure S1:**
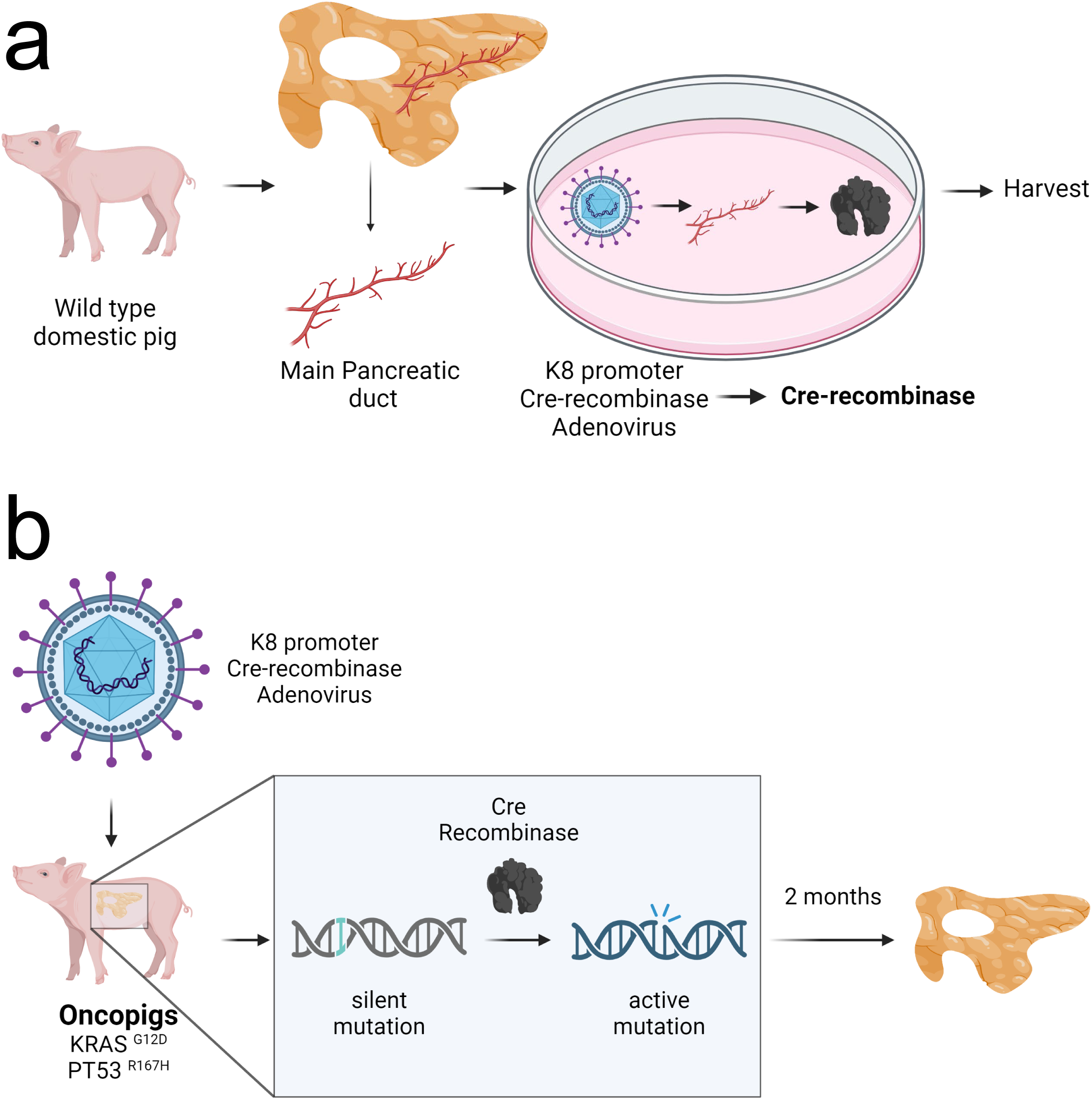
Schematic Overview of Ad-K8-Cre Adenovirus Experimental Procedure. (a) *Ex Vivo* Assessment in Wild Type Porcine Pancreas: This panel illustrates the workflow starting from a wild type domestic pig, extraction of the main pancreatic duct, and subsequent ex vivo infection with Ad-K8-Cre adenovirus. The adenovirus, driven by the Keratin 8 promoter to express Cre-recombinase, targets pancreatic epithelial cells cultured in a petri dish, followed by harvesting for analysis. (b) *In Vivo* targeted Activity in Oncopigs: This diagram outlines the in vivo process where the Oncopig, harboring latent KRAS^G12D^ and TP53^R167H^ mutations, is injected with the adenovirus directly into the pancreatic duct. The illustration shows the mechanism by which Cre-recombinase activates these mutations, leading to observable changes in the pancreas after 2 months.

**Figure S2:**
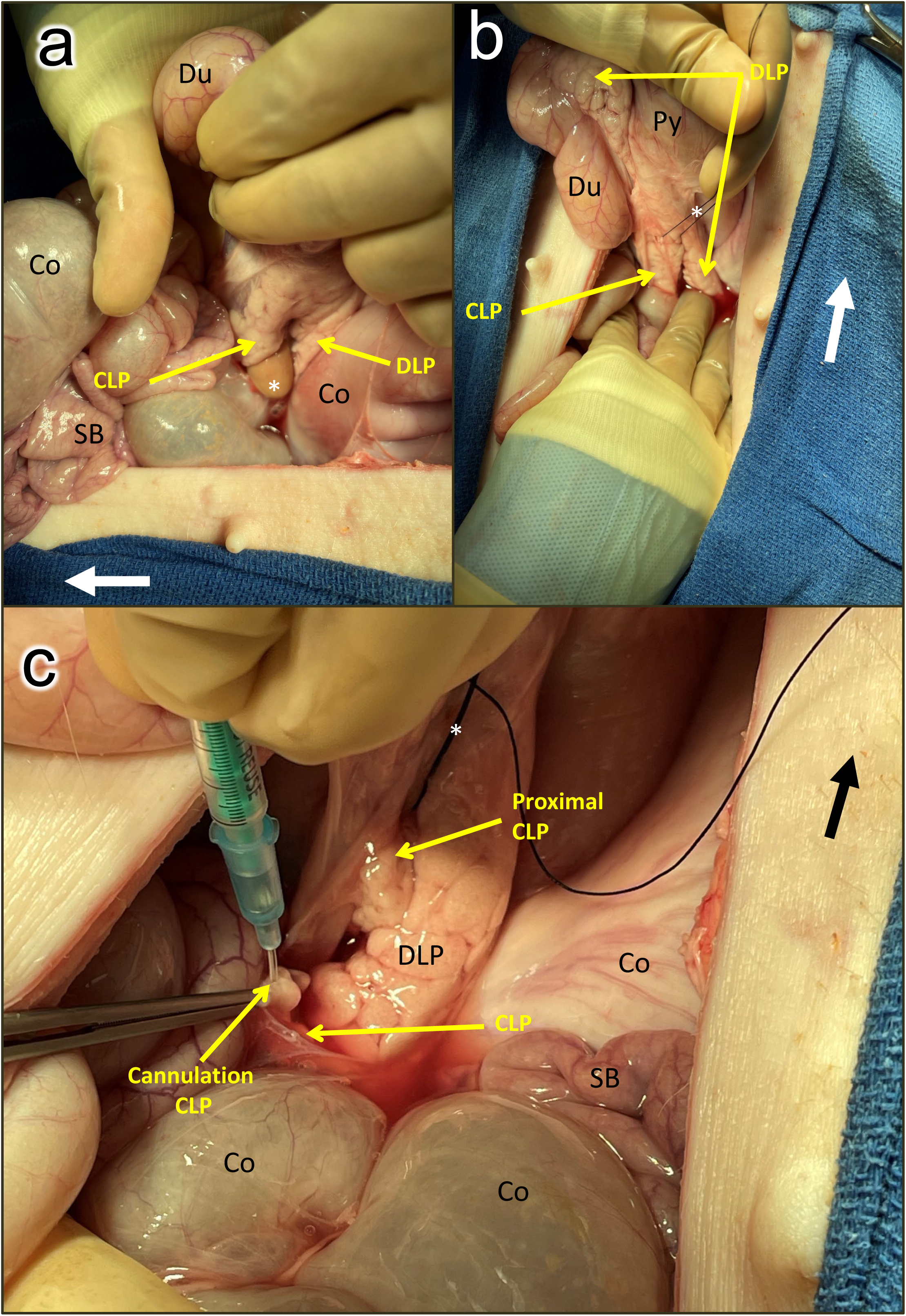
Effect of Ad-K8-Cre injection on gross pancreatic morphology in the Oncopig. (a) Freshly excised connecting lobe of the pancreas from an Oncopig injected with a high dose (10^9^ pfu) of Ad-K8-Cre, showing atrophy and nodularity in the region of the injection (dashed circle). Thin arrow = region of ductal cannulation for vector injection; dashed oval = region of gross effect; thick arrow = cephalad; ruler = in/ cm. (b) Same as panel a, except with a low dose (10^7^ pfu) of Ad-K8-Cre. (c) Regional *in vivo* appearance (necropsy) in Oncopig with a high dose injection. The midline incision has been re-opened and the superior margin of the pancreas has been exposed. SB = small bowel; thick arrow = cephalad. (d) Same as panel c, but the duodenal lobe of the pancreas (DLP) has been reflected cephalad to expose the connecting lobe of the pancreas (CLP, the region of injection). Co = colon; Du = duodenum; Py = pylorus; St = stomach.

**Figure S3:**
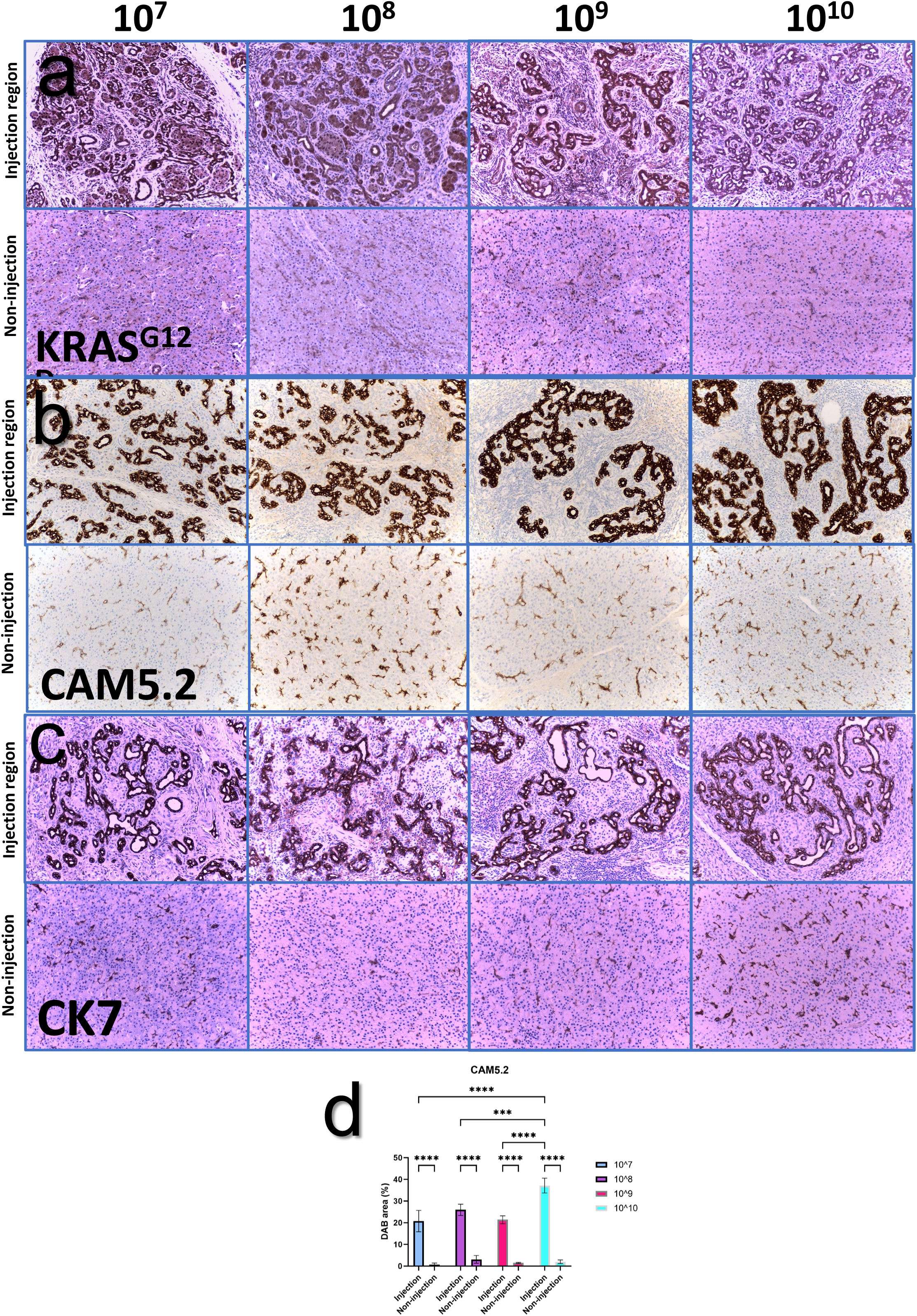
Expression of Oncogenic KRAS and Epithelial Markers in Pancreatic Tissues Post Ad-K8-Cre Injection. (a) KRAS^G12D^ Staining: Immunohistochemical analysis of KRAS^G12D^ in pancreatic tissue sections from Oncopigs injected with Ad-K8-Cre (10^7^ to 10^10^ pfu). The top row represents injection site regions, showing a dose-dependent increase in KRAS^G12D^ expression within ductal epithelial cells. The bottom row displays non-injection sites, with no significant KRAS^G12D^ staining as internal controls. (b) CAM5.2 Staining: CAM5.2 immunostaining demonstrates epithelial marker expression in ductal cells at the injection site across varying Ad-K8-Cre doses, confirming the ductal localization of the viral vector. Non-injection site regions (bottom row) reveal minimal CAM5.2 expression. (c) CK7 Staining: CK7 immunohistochemistry shows additional epithelial marker expression, corroborating the localization of Ad-K8-Cre activity in ductal structures. Higher doses (10^9^ and 10^10^ pfu) exhibit intensified CK7 labeling at injection sites, while non-injection sites maintain baseline CK7 expression. (d) Quantification of CAM5.2 Expression: Bar graph quantifies DAB-positive area for CAM5.2 across different doses at injection and non-injection sites. Statistical significance is noted as follows: ****p<0.0001, ***p<0.001, **p<0.01, *p<0.05.

**Figure S4:**
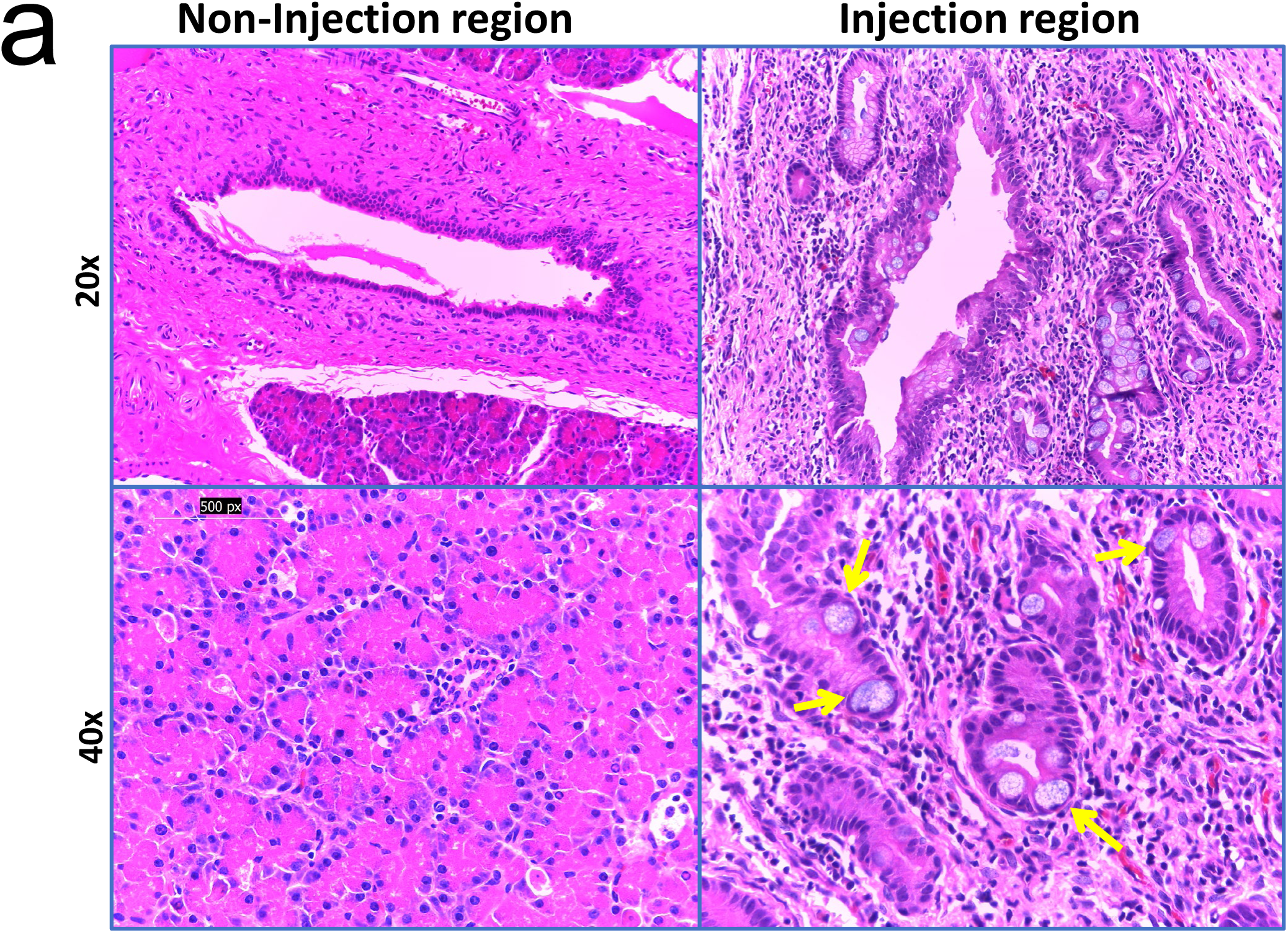
Histopathological Analysis and Genotyping of Oncopigs Post Ad-K8-Cre Injection. (a) Hematoxylin and Eosin (H&E) Staining in Injection and Non-Injection Regions: Representative H&E-stained sections at 20x (top row) and 40x (bottom row) magnifications. Left panels show non-injection regions with preserved normal pancreatic architecture. Right panels display injection regions, with yellow arrows indicating Goblet cells. (b) Genotyping of Oncopigs for Verification of KRAS^G12D^ Mutation: Gel electrophoresis results confirm the presence of the KRAS^G12D^ mutation in genetically engineered Oncopigs. Lanes represent individual samples, with positive and negative controls on the left. The presence of a specific amplicon at the expected size validates successful genotype targeting in Oncopigs used for the study.

**Figure S5:**
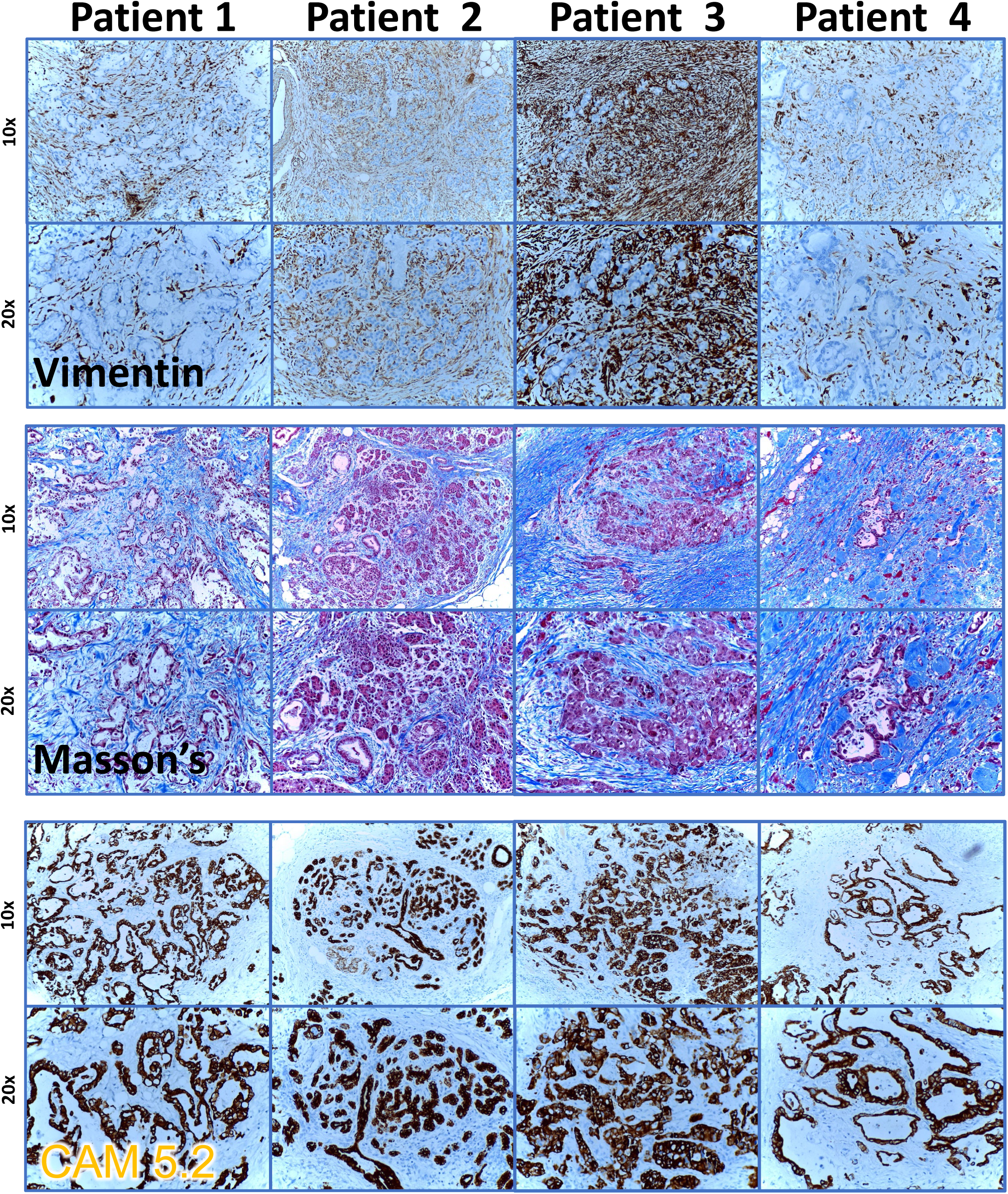
Comparative Histopathological Analysis of Human Pancreatic Cancer Samples. Vimentin Staining (Top Two Rows): Showcases vimentin immunohistochemistry across PDAC samples from four patients, highlighting the stromal reaction characteristic of pancreatic cancer. Vimentin is abundantly expressed, indicating a robust stromal response. Higher magnifications (20x) provide a closer view of the stromal involvement within the cancer architecture. Masson’s Trichrome Staining (Middle Two Rows): Depicts collagen deposition within the tumor microenvironment, stained in blue. These sections reveal varying levels of fibrosis across different samples, with increased collagen indicative of a desmoplastic reaction, which is a hallmark of pancreatic cancer. CAM5.2 Staining (Bottom Two Rows): Highlights the expression of cytokeratin in epithelial cells, underscoring the epithelial nature of the cancerous cells.

